# Simplicity: web-based visualization and analysis of high-throughput cancer cell line screens

**DOI:** 10.1101/2023.09.08.556619

**Authors:** Alexander L. Ling, Weijie Zhang, Adam Lee, Yunong Xia, Mei-Chi Su, Robert F. Gruener, Sampreeti Jena, Yingbo Huang, Siddhika Pareek, Yuting Shan, R. Stephanie Huang

## Abstract

High-throughput drug screens are a powerful tool for cancer drug development. However, the results of such screens are often made available only as raw data, which is intractable for researchers without informatic skills, or as highly processed summary statistics, which can lack essential information for translating screening results into clinically meaningful discoveries. To improve the usability of these datasets, we developed Simplicity, a robust and user-friendly web interface for visualizing, exploring, and summarizing raw and processed data from high-throughput drug screens. Importantly, Simplicity allows for easy recalculation of summary statistics at user-defined drug concentrations. This allows Simplicity’s outputs to be used with methods that rely on statistics being calculated at clinically relevant doses. Simplicity can be freely accessed at https://oncotherapyinformatics.org/simplicity/.

## 1 Introduction

In the past decade, multiple institutions have generated publicly available datasets for hundreds of compounds screened in hundreds of cancer cell lines (CCLs) (Ling *et al*., 2018). Substantial efforts have been made to harmonize and distribute data from these datasets both via programmatic (Smirnov *et al*., 2016) and web-based (Tsherniak *et al*., 2017; Smirnov *et al*., 2018) interfaces. However, programmatic access is challenging for researchers who lack coding or bioinformatic experience, and web-based interfaces for these datasets do not currently provide users with the means to summarize drug efficacy at specific drug concentrations or concentration ranges.

Given recent evidence that CCL screening data should be analyzed at clinically achievable drug concentrations to generate clinically relevant findings (Ling and Huang, 2020) and the recent deployment of a web-based interface for utilizing CCL screening data to predict drug combination efficacy in a dose-dependent fashion (Yunong Xia *et al*., 2023), we developed the Simplicity (<u>S</u>implified <u>I</u>nterface to <u>M</u>anipulate <u>P</u>rec<u>L</u>inical <u>I</u>nformation for <u>C</u>ancer <u>I</u>n vitro <u>T</u>herap<u>Y</u>) web-interface to enable researchers without programming experience to easily perform dose-dependent calculations with CCL screening data.

## 2 Methods and software implementation

Raw screening data was obtained from four large CCL screening datasets:

1. The Cancer Therapeutics Response Portal v2 (CTRPv2) (Basu *et al*., 2013; Seashore-Ludlow *et al*., 2015; Rees *et al*., 2016)
2&3. Genomics of Drug Sensitivity in Cancer 1 & 2 (GDSC1 and GDSC2) (Iorio *et al*., 2016; Yang *et al*., 2013; Garnett *et al*., 2012)
4. PRISM Repurposing (Corsello *et al*., 2020)

CTRPv2 was generated at the Broad Institute between 2012 and 2013 and contains data for 544 compounds screened in 887 cell lines. GDSC1 was generated by Massachusetts General Hospital and the Wellcome Sanger Institute between 2010 and 2015 and contains data for 343 compounds screened in 987 cell lines, with a follow up screen (GDSC2) being performed by Sanger between 2015 and 2017 for 192 compounds in 809 cell lines. PRISM Repurposing was published by the Broad Institute in 2020 and contains screening data for 1446 compounds in 481 cell lines. Further details for these screens can be found in the “Data Explorer/Explore Datasets” tab of Simplicity or in their respective publications.

Full details of how these datasets were harmonized and quality controlled are included in the **Supplemental Methods**. However, a very brief description of this process is as follows.

Initial cell line and compound harmonization tables were taken from our prior harmonization efforts (Ling *et al*., 2018; Ling and Huang, 2020), which included harmonized cell line and compound IDs for CTRPv2 and GDSC1. Data was further harmonized and annotated using a mix of manual curation as well as data from https://www.cellosaurus.org/, the BROAD Drug Repurposing Hub (https://www.broadinstitute.org/drug-repurposing-hub), and https://webchem.org/. Raw data from each dataset was then quality controlled, and dose-response curves were fit to the harmonized and quality controlled data. A user interface for exploring and manipulating this data was created using the shiny package (Winston Chang *et al*., 2020) in R (R Core Team, 2020). This interface, Simplicity, was then deployed on scalable cloud-based infrastructure.

## 3 Validation of data quality

To validate the quality of Simplicity’s refitted dose-response curves, cross-dataset agreement was measured for shared compounds and cell lines under the hypothesis that compound/cell-line pairs which were screened in multiple screens should result in similar AUC values across the same dose-range in both screens. As such, high correlation in drug sensitivities measured between two screens should indicate that dose-response curves have been appropriately fit, while lower correlations may indicate inferior curve-fitting approaches.

We took data from three sources of harmonized data for the drug screens included in Simplicity and sought to ensure that the cross-dataset agreement in Simplicity was not inferior to other available sources. These three sources were:

1. Simplicity
2. Corsello *et al*., 2020
3. PharmacoGx (Smirnov *et al*., 2016)

Cross-dataset correlations were similar between all datasets when using any of the three data sources, with larger variations between sources noted when comparing drug sensitivities measured in PRISM-Repurposing to other screens (**Figures S1-S3**). Despite similar performance between data sources, a few compounds were much more or less correlated between screens with Simplicity than with other datasets.

To understand these situations, we plotted PRISM-Repurposing vs. CTRPv2 AUC values for the top eight compounds in which PharmacoGx had higher cross-dataset correlations than Simplicity (**Figure S4**) and the top eight compounds in which Simplicity had higher cross-dataset correlations than PharmacoGx (**Figure S5**). This data suggests that the majority of compounds that see large differences in Spearman’s rho values between data sources are compounds that have low efficacies in most tested cell lines, resulting in relatively little variation in measured drug sensitivities. While it does appear that the curve fitting approach used by Simplicity may perform worse or better for specific compounds than the approaches used by other data sources, average performance across all tested compounds is very similar.

This gives us confidence that the new functionalities provided by Simplicity to non-computational users of these datasets do not come at a cost of reduced data quality. These functionalities are described in the following sections.

## 4 Visualizing screening data with Simplicity

Simplicity allows users to generate customized plots to easily visualize information such as:

1. Ancestry (**Figure 1A**), age, gender, and cancer types across specific CCL populations (not shown). This can facilitate rapid intuition around how well a set of CCLs represents a researcher’s patient cohort of interest.
2. Summary statistics of drug sensitivity across many CCLs for a single drug or across many drugs for a single CCL (**Figure 1B**). This enables users to quickly identify which cell lines are most or least sensitive to a given drug or to identify which drugs a given cell line shows exceptional sensitivity/resistance to.
3. Raw data for a given drug/CCL pair’s dose-response curve (**Figure 1C**). This allows users to directly visualize the quality of a given dose-response curve, as well as to determine the level of reproducibility for a given drug/CCL pair across different datasets and replicates.
4. Relevant background information to the results being plotted, such as information about variations in assay conditions between different CCLs screens and different experimental runs within a given screen (**Figure 1D**). This can allow users to easily visualize how factors such as cell seeding density, plate format, assay reagent, and treatment duration influence dose-response curves.

Customization of these plots is achieved via use of searchable drop-down menus and slider bars which allow filtering based on such characteristics as CCL disease type, age, gender, and ancestry makeup or compound molecular target, mechanism of action, or clinical phase.

**Figure 1.**
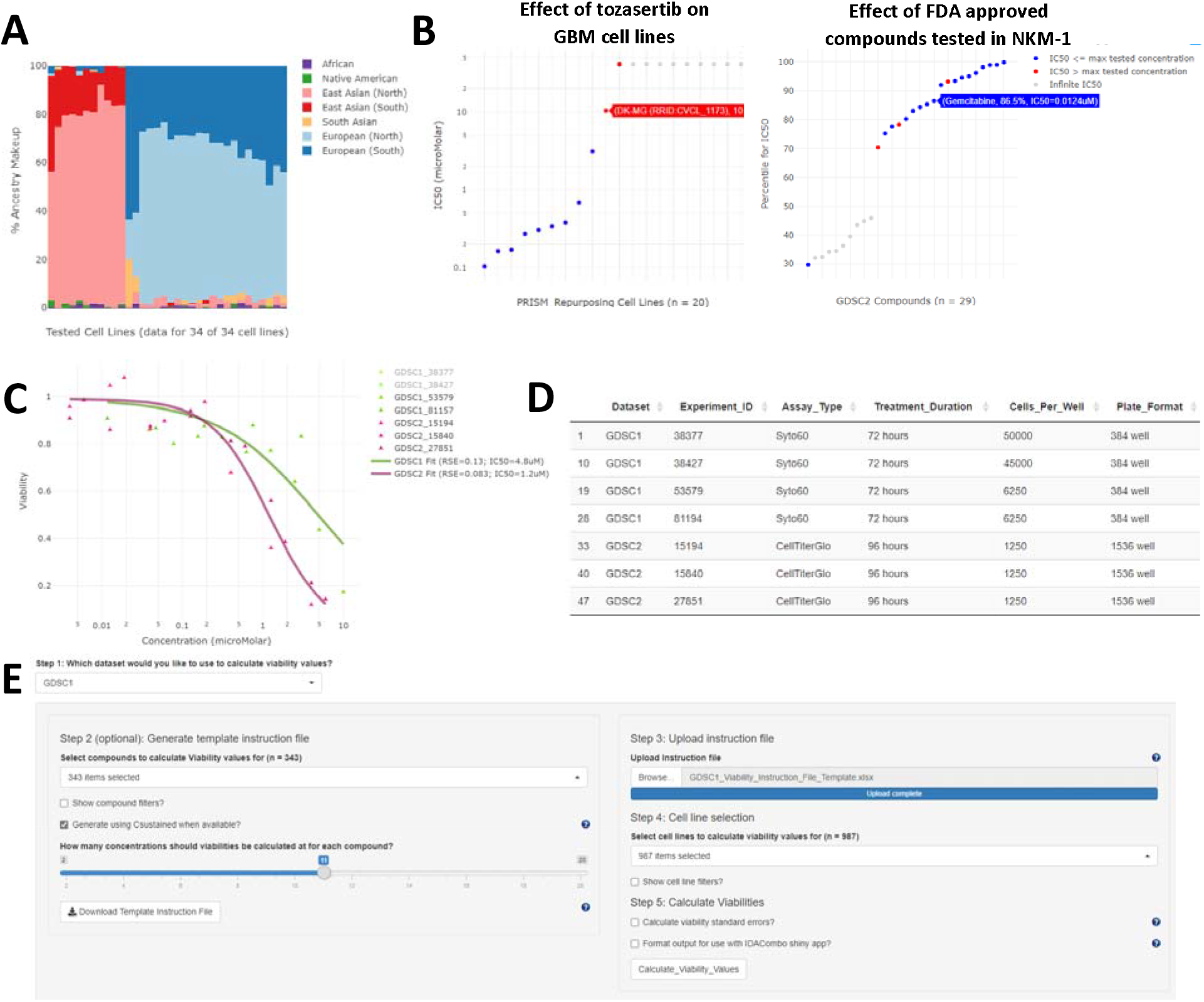
Example functionality of Simplicity. Plots, tables, and interfaces from Simplicity. **(A)** Ancestry plot for glioblastoma (GBM) cell lines tested with 5-Fluorouracil in GDSC1 as provided by the “Data Explorer/Explore Compounds” tab. **(B)** Examples of drug and cell-line level summaries produced by Simplicity. <u>Left panel</u>: Plot showing measured sensitivities (IC50s) of Tozasertib in GBM cell lines in the PRISM-Repurposing dataset as provided by the “Data Explorer/Explore Compounds” tab. Cell lines names and exact IC50 values can be obtained by hovering over each data point. <u>Right panel</u>: Plot showing relative sensitivity of NKM-1 cell line to FDA approved (Launched) compounds tested in GDSC2 as measured by IC50 percentile relative to all other cell lines tested with each compound in GDSC2 as provided by the “Data Explorer/Explore Cell Lines” tab. Higher percentiles indicate NKM-1 was more sensitive to a given compound relative to other tested lines. Direct IC50 values can be obtained by hovering over each data point or by downloading the summary statistics tables provided in the “Download Bulk Data” tab of Simplicity. Note that infinite IC_50_ values occur when fitted dose-response curves have a lower asymptote above 50% viability. This can occur when the data directly implies an asymptote above 50% viability or when the tested compound shows no efficacy at any tested dose such that the fitted dose response curve is simply a flat line at 100% viability. **(C)** Calculated dose-response curves for cisplatin in the NKM-1 cell line in both GDSC1 and GDSC2 along with the experiment IDs used to calculate the curves as provided by the “Data Explorer/Plot Dose-Response Curves” tab. **(D)** Table of experimental conditions used in the experiments shown in panel C as provided by the “Data Explorer/Plot Dose-Response Curves” tab. **(E)** User interface for calculating viability values at specified concentrations. The interface allows users to easily select compounds, cell lines, and concentrations of interest using a graphical user interface. A similar interface is also available for calculating area under the curve (AUC) values at custom concentration ranges.

## 5 Calculating custom summary statistics with Simplicity

To enable researchers to easily generate dose-specific metrics of drug efficacy from these screens, Simplicity provides the “Calculate Custom Statistics/AUC Values” and “Calculate Custom Statistics/Viability Values” tabs to calculate AUC and Viability values at custom concentrations/concentration ranges using a simple graphical user interface (**Figure 1E**). The interface provides the same searchable drop-down menus and slider bars present throughout the rest of the app to allow easy selection of compounds and CCLs of interest. The results of these calculations are provided as downloadable tables, with an option to automatically format the output for direct use with the IDACombo web application, which uses dose-specific estimates of monotherapy drug efficacy to predict drug combination efficacy across different doses of combined drugs (Yunong Xia *et al*., 2023).

## 6 Accessing bulk data through Simplicity

Simplicity also provides bulk data download for researchers who wish to use Simplicity’s harmonized data with their own informatic tools. These can be accessed via the “Download Bulk Data” tab. Available data includes:

1. Harmonized CCL and compound names between the included datasets.
2. Clinically relevant concentrations for 143 clinically tested compounds that are included in Simplicity.
3. AUC and IC_50_ values for the CCL-compound pairs tested in each screen.
4. Raw viability values from each screen following compound and CCL name harmonization.

## 7 Summary

Simplicity provides a graphical user web interface which allows users to easily visualize and manipulate data from high-throughput CCL drug screens. Notably, Simplicity provides the ability to query viability and AUC values at custom doses/dose ranges, enabling analyses to be conducted with clinically relevant concentrations without the need for coding or informatic experience. It is our hope that this will remove a significant barrier for non-computational scientists who wish to use these datasets to conduct such dose-dependent studies. A video tutorial on the use of Simplicity is available at https://www.youtube.com/watch?v=oNuwRDs_5DQ.

## Author Contributions

Conceptualization and App Development: ALL, RSH

App Beta Testing: ALL, WZ, AL, YX, MCS, RG, SJ, YH, SP, YS

App Maintenance: ALL, WZ, AL, RSH

Manuscript Writing: ALL

Manuscript Review and Editing: ALL, WZ, MCS, RSH

