## Supplemental Materials for "Simplicity: web-based visualization and analysis of high-throughput cancer cell line screens"

### Supplemental Methods

#### Dataset sources

Data for the Cancer Therapeutics Response Portal version 2 (CTRPv2)^1–3^ was obtained from the following files:

- *CTRPv2.0_2015_ctd2_ExpandedDataset.zip*: downloaded from <https://ctd2-data.nci.nih.gov/Public/Broad/CTRPv2.0_2015_ctd2_ExpandedDataset/CTRPv2.0_2015_ctd2_ExpandedDataset.zip> on 5/29/2020.
- *CTRPv2.0._COLUMNS.xlsx*: downloaded from <https://ctd2-data.nci.nih.gov/Public/Broad/CTRPv2.0_2015_ctd2_ExpandedDataset/CTRPv2.0._COLUMNS.xlsx> on 5/29/2020.
- *CTRPv2.0._INFORMER_SET.xlsx*: downloaded from <https://ctd2-data.nci.nih.gov/Public/Broad/CTRPv2.0_2015_ctd2_ExpandedDataset/CTRPv2.0._INFORMER_SET.xlsx> on 5/29/2020.

Data for the Genomics of Drug Sensitivity in Cancer 1 and 2 (GDSC1 and GDSC2)^4–6^ was obtained from the following files:

- *GDSC_Raw_Data_Description.pdf*: downloaded from <ftp://ftp.sanger.ac.uk/pub4/cancerrxgene/releases/release-8.2/GDSC_Raw_Data_Description.pdf> on 4/30/2020
- *GDSC1_public_raw_data_25Feb20.csv*: downloaded from [ftp://ftp.sanger.ac.uk/pub4/cancerrxgene/releases/release-8.2/GDSC1_public_raw_data_25Feb20.csv](ftp://ftp.sanger.ac.uk/pub4/cancerrxgene/releases/release-8.2/GDSC1_public_raw_data_25Feb20.csv%20) on 4/30/2020
- *GDSC2_public_raw_data_25Feb20.csv*: downloaded from <ftp://ftp.sanger.ac.uk/pub4/cancerrxgene/releases/release-8.2/GDSC2_public_raw_data_25Feb20.csv> on 4/30/2020
- *screened_compounds_rel_8.2.csv*: downloaded from <ftp://ftp.sanger.ac.uk/pub4/cancerrxgene/releases/release-8.2/screened_compunds_rel_8.2.csv> on 5/14/2020
- *Cell_Lines_Details.xlsx*: downloaded from <ftp://ftp.sanger.ac.uk/pub4/cancerrxgene/releases/release-8.2/Cell_Lines_Details.xlsx> on 5/14/2020
- *GDSC1_fitted_dose_response_25Feb20.csv*: downloaded from <ftp://ftp.sanger.ac.uk/pub4/cancerrxgene/releases/release-8.2/GDSC1_fitted_dose_response_25Feb20.csv> on 6/9/2020
- *GDSC2_fitted_dose_response_25Feb20.csv*: downloaded from <ftp://ftp.sanger.ac.uk/pub4/cancerrxgene/releases/release-8.2/GDSC2_fitted_dose_response_25Feb20.csv> on 6/9/2020
- *GDSC_Fitted_Data_Description.pdf*: downloaded from [ftp://ftp.sanger.ac.uk/pub4/cancerrxgene/releases/release-8.2/GDSC_Fitted_Data_Description.pdf on 6/9/2020](ftp://ftp.sanger.ac.uk/pub4/cancerrxgene/releases/release-8.2/GDSC_Fitted_Data_Description.pdf%20on%206/9/2020)

Data for the PRISM Drug Repurposing Resource (PRISM-Repurposing)^7^ was obtained from the following files:

- *secondary-screen-logfold-change.csv*: downloaded from <https://ndownloader.figshare.com/files/20237730> on 4/6/2020
- *secondary-screen-mfi.csv*: downloaded from [https://ndownloader.figshare.com/files/20237742](https://ndownloader.figshare.com/files/20237742%20) on 4/6/2020
- *secondary-screen-cell-line-info.csv*: downloaded from <https://ndownloader.figshare.com/files/20237769> on 4/6/2020
- *secondary-screen-replicate-treatment-info.csv*: downloaded from <https://ndownloader.figshare.com/files/20237760> on 4/6/2020
- *secondary-screen-pooling-info.csv*: downloaded from <https://ndownloader.figshare.com/files/20237766> on 4/6/2020
- *sample_info.csv*: downloaded from <https://ndownloader.figshare.com/files/21522000> on 4/10/2020
- *repurposing_samples_20180907.txt*: downloaded from [https://clue.io/data/REP#](https://clue.io/data/REP) on 4/10/2020
- *secondary-screen-dose-response-curve-parameters.csv*: downloaded from <https://ndownloader.figshare.com/files/20237739> on 4/14/2020

The following cell line and compound harmonization files were obtained from Ling et al., 2018^8^:

- *Table S2_Screened Cell Line Info_Ling et al_2018.xlsx*
- *Table S3_Screened Drug Info_Ling et al_2018.xlsx*

Additional cell line information, including ancestry information, was obtained from https://www.cellosaurus.org/:

- *cellosaurus.xml*: downloaded from [ftp://ftp.expasy.org/databases/cellosaurus/cellosaurus.xml on 5/29/2020](ftp://ftp.expasy.org/databases/cellosaurus/cellosaurus.xml%20on%205/29/2020).

Additional compound information, including clinical trial phase, was obtained from the BROAD Drug Repurposing Hub:

- *repurposing_drugs_20200324*: downloaded from <https://s3.amazonaws.com/data.clue.io/repurposing/downloads/repurposing_drugs_20200324.txt> on 5/29/2020.

#### Pre-processing raw data from PRISM-Repurposing

Given that the PRISM Repurposing screen tested cell lines as pooled groups, we developed a cleaning strategy which leverages information from all cell lines in a pool when scaling assay values. This cleaning strategy is as follows:

1. Cell lines which failed short-tandem-repeat (STR) fingerprinting for authentication were removed from the analysis.
2. For each plate that was screened, vehicle control barcode values were scaled as follows:
   a. All barcodes quantified in each vehicle control well were scaled by dividing the geometric mean of the control barcodes in that well (control-gmean) by the median control-gmean across all vehicle control wells on that plate.
   b. Any wells which had a control-gmean that was <5% of the median control-gmean on the plate (i.e. control wells where signal was very low relative to other control wells, as might be expected from a significant pipetting error) were discarded from the analysis.
3. The effect of tested drugs on each cell line tested on a plate were calculated as follows:
   a. The median fluorescence intensity (MFI) value for each barcode in a given well was scaled by dividing it by the geometric mean of the spike-in barcode controls in that well.
   b. Viability values for a given barcode in a given well were calculated by dividing the scaled MFI values for that barcode by the geometric mean of the scaled vehicle control well MFI values for that barcode on that plate.
   c. A 4-parameter log-logistic curve was fit to each cell line-drug pair on the plate (see Fitting dose-response curves section), and the residual distance between measured and fitted viabilities were calculated for each drug dose. A multiplication factor was then calculated for each cell line-drug-dose trio which would match measured viabilities to fitted viabilities. Under the assumption that these multiplication factors should be normally distributed around 1 across all cell lines pooled in a single well and would only produce a distribution with a mean not equal to 1 if a systematic error existed in that well, we then took the geometric mean of all multiplication factors calculated for each well and applied that mean multiplication factor to viability values for all cell lines in that well. Note that this correction was only performed for wells in which at least 3 cell lines could be successfully fit to a 4-parameter log-logistic curve with the other data on the plate.
4. For each detection pool-screen pair
   a. We corrected data for drift by analyte id on a per detection pool-screen pair basis. Our correction method assumes that the large majority of tested drugs on any given detection plate will result in 100% viability in a given cell line (analyte_id) at concentrations less than 1 nM. This correction uniformly shifts viabilities across all drugs and doses for a given analyte_id-plate pair by shifting median viabilities for drug concentrations < 1 nM to be 100%.

#### Fitting drug dose-response curves

1. For each screen:
   a. A pre-fit model was created for each drug/cell-line pair by fitting data to a 4-parameter log logistic curve under the following constraints:
      i. 10 <= slope of dose response curve (parameter b) >= 0
      ii. 0.99 >= lower asymptote (parameter c) >= 0
      iii. 1.01 >= upper asymptote (parameter d) >= 0.99
      iv. ED50 (parameter e) >= minimum non-zero dose tested / 2
   b. If the fit was successful and had a residual standard error (RSE) <= 0.4 (for PRISM-Repurposing) or <= 0.2 (for CTRPv2, GDSC1, and GDSC2), that drug/cell-line pair was retained in that screen. If not, that pair was discarded from the screen.
2. All retained data was used to fit 4-parameter log-logistic curves under the same constraints now using data across all retained screens for each drug/cell-line pair.
   a. If the resulting fit was successful and had an RSE <= 0.4 (for PRISM-Repurposing) or <= 0.2 (for CTRPv2, GDSC1, and GDSC2), the fit for that drug/cell-line pair was considered to be of high enough quality to be included in Simplicity.

**Calculating Area Under the Curve (AUC) values**

AUC values are calculated by taking integrals of 4-parameter log-logistic curves in which natural logs are used with all concentration values.

#### Comparing Simplicity to other drug screening resources

*PharmacoGx based overlap:*

The PharmacoGx package (v3.2.0)^9^ was used to download the following four datasets (PSets) on 11/11/2022: 1. GDSC2_2020(v1-8.2); 2. GDSC_2020(v2-8.2); 3. CTRPv2_2015; 4. PRISM_2020.

Cell lines were matched to Simplicity data using the “sampleid” field of each PSet with a manually curated map of Simplicity cell line names to PharmacoGx sample IDs (**Supplementary Table S1**). Note that we omitted PharmacoGx experiments with the “Lu-99A” and “COLO 320HSR” sample IDs from our screen overlap analysis as there is reason to believe that multiple versions of these cell lines were used by different datasets and may not be directly comparable.

Compounds were matched to Simplicity data using the “treatmentid” field of each PSet with a manually curated map of Simplicity compound names to PharmacoGx treatment IDs (**Supplementary Table S2**). Note that we omitted blebbistatin, bleomycin A2, and nutlin-3 from overlap analyses between some screens (CTRPv2/PRISM vs GDSC1/GDSC2) as different chemical forms of these compounds were used in different screens which may not be directly comparable.

AUC values for PharmacoGx were calculated using recomputed Hill Curve coefficients from PharmacoGx with the computeAUC() function provided by the package and restricting calculations to the shared concentration range between compared datasets.

*Corsello et al. based overlap:*

Comparisons of PRISM vs CTRPv2 and GDSC1 from Corsello et al.^7^ were downloaded on 9/23/2020 from: <https://static-content.springer.com/esm/art%3A10.1038%2Fs43018-019-0018-6/MediaObjects/43018_2019_18_MOESM2_ESM.xlsx>. 4-parameter log-logistic coefficients provided in the “11-Reprocessed AUCs” sheet of the workbook were used to calculate AUC values across shared concentration ranges between datasets. Cell lines and compounds were matched between this dataset and Simplicity using the original cell line and compound names from each dataset with the harmonization tables provided in the “Download Bulk Data” tab of Simplicity (<https://oncotherapyinformatics.org/simplicity/>).

*Simplicity based overlap:*

Recalculated 4-parameter log-logistic parameters were used to recalculate AUC values for shared concentration ranges between different datasets. Cell line and compound overlap were determined based on the harmonization tables provided in the “Download Bulk Data” tab of Simplicity (<https://oncotherapyinformatics.org/simplicity/>).

*Comparisons of overlap between different dataset sources:*

Spearman’s rho values were estimated with 95% confidence intervals between AUC values for each given overlapping compound using only overlapping cell lines and concentration ranges for each dataset.

**Software**

Dose-response curve fitting and data preparation for the Simplicity app was performed using R (v3.6.1)^10^ with the following packages: drc (v3.0.1)^11^, progress (v1.2.2)^12^, doSNOW (v1.0.18)^13^, readr (v1.3.1)^14^, readxl (v1.3.1)^15^, openxlsx (v4.1.4)^16^, XLM (v4.0-0)^17^, webchem (v1.0.0)^18^, and Hmisc (v4.4-0)^19^.

Comparison of Simplicity to other repositories of high-throughput screening data was performed using R (v4.2.1)^20^ with the following packages: PharmacoGx package (v3.2.0)^9^, DescTools (0.99.47)^21^, and readxl (v1.4.1)^22^.

The Simplicity app itself is run using R (v4.0.3)^23^ with the following packages: shiny (v1.5.0)^24^, shinyhelper (v0.3.2)^25^, shinyWidgets (v0.5.4)^26^, shinybusy (v0.2.2)^27^, readr (v1.4.0)^28^, readxl (v1.3.1)^15^, plotly (v4.9.2.2)^29^, ggplot2 (v3.3.3)^30^, openxlsx (v4.2.3)^31^, drc (v3.0-1)^11^, ipc (v0.1.3)^32^, promises (v1.1.1)^33^, future (v1.21.0)^34^, doFuture (v0.12.2)^34^, memuse (v4.1-0)^35,36^, DT (v0.16)^37^, and tools (included with R v4.0.3)^23^.

**Web hosting**

Simplicity is hosted on virtual machines (VMs) purchased from DigitalOcean (<https://www.digitalocean.com/>). Each VM is running Ubuntu 20.04.1 LTS (GNU/Linux 5.4.0-51-generic x86_64) with 8 virtual CPUs, 16 GB RAM, and 100 GB disk space. Traffic is split across VMs using a load balancer. Connections within each VM are handled via Apache server 2.4 and shiny server v1.5.15.953-amd64.

**Data accessibility**

Compiled and raw data used by Simplicity is available for download from the website itself (<https://oncotherapyinformatics.org/simplicity/>) and from the projects OSF repository (<https://osf.io/a9w5r/>). Code for the Simplicity app is available at <https://github.com/Alexander-Ling/Simplicity>.

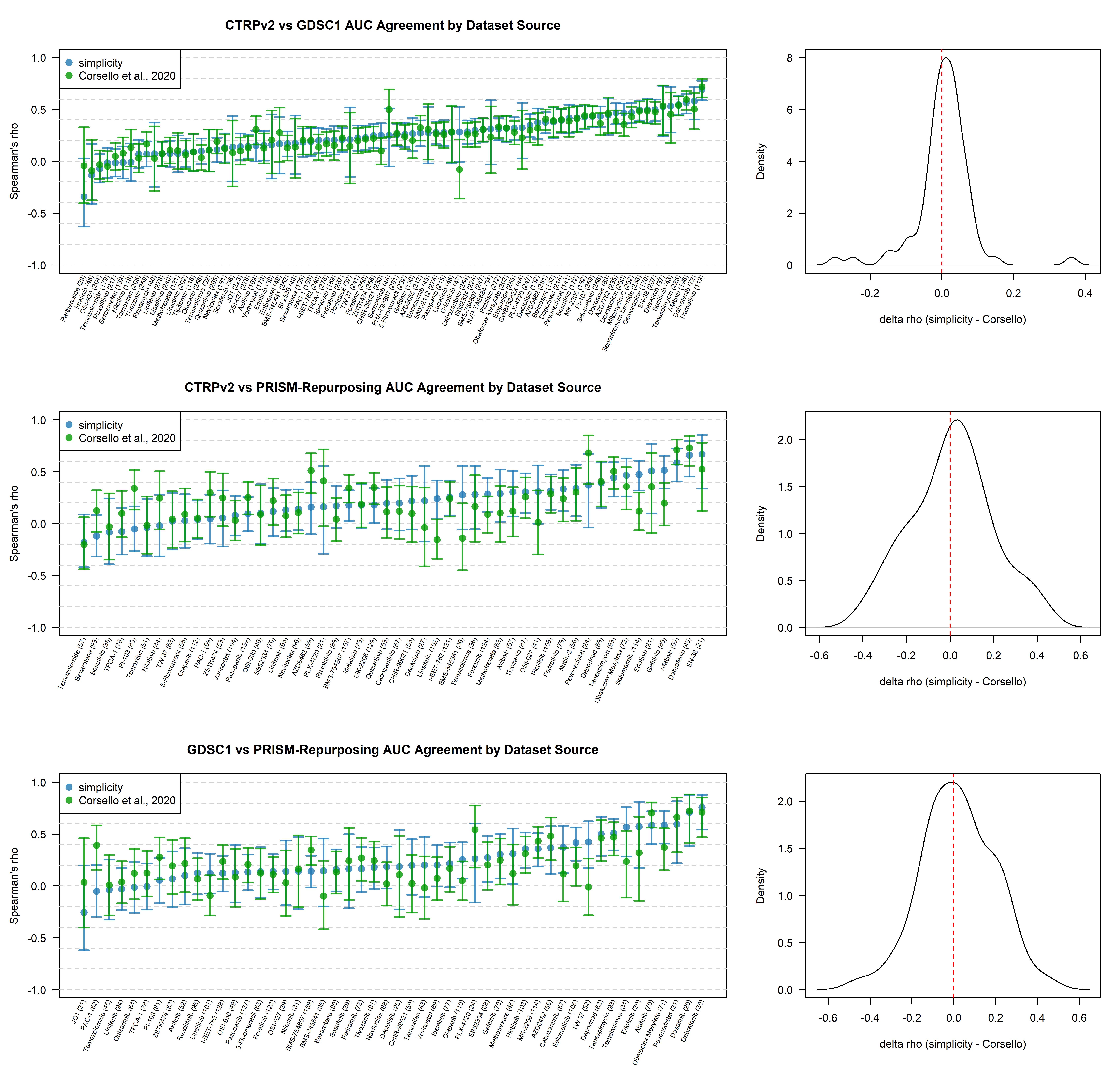
**Figure S1. Agreement between screening datasets using data from Simplicity vs. from Corsello et al, 2020**^7^**.** Agreement between three dataset comparisons is shown: 1. (**A**) CTRPv2 vs. GDSC1; 2. (**B**) CTRPv2 vs. PRISM-Repurposing; 3. (**C**) GDSC1 vs PRISM-Repurposing. Agreement between datasets is quantified as Spearman’s rho for AUC values calculated for each compound using overlapping cell lines and drug concentrations. Plots on the **LEFT** show these rho values for each drug with error bars indicating 95% confidence intervals. Density plots on the **RIGHT** show the distribution of differences in rho values between Simplicity and Corsello et al., with negative values indicating higher rho values using data from Corsello et al. and positive values indicating higher rho values using Simplicity. Parentheses after compound names indicate the number of shared cell lines between the two datasets. Only compounds tested in at least 20 shared cell lines between the datasets were included in the analysis.

**C)**

**B)**

**A)**

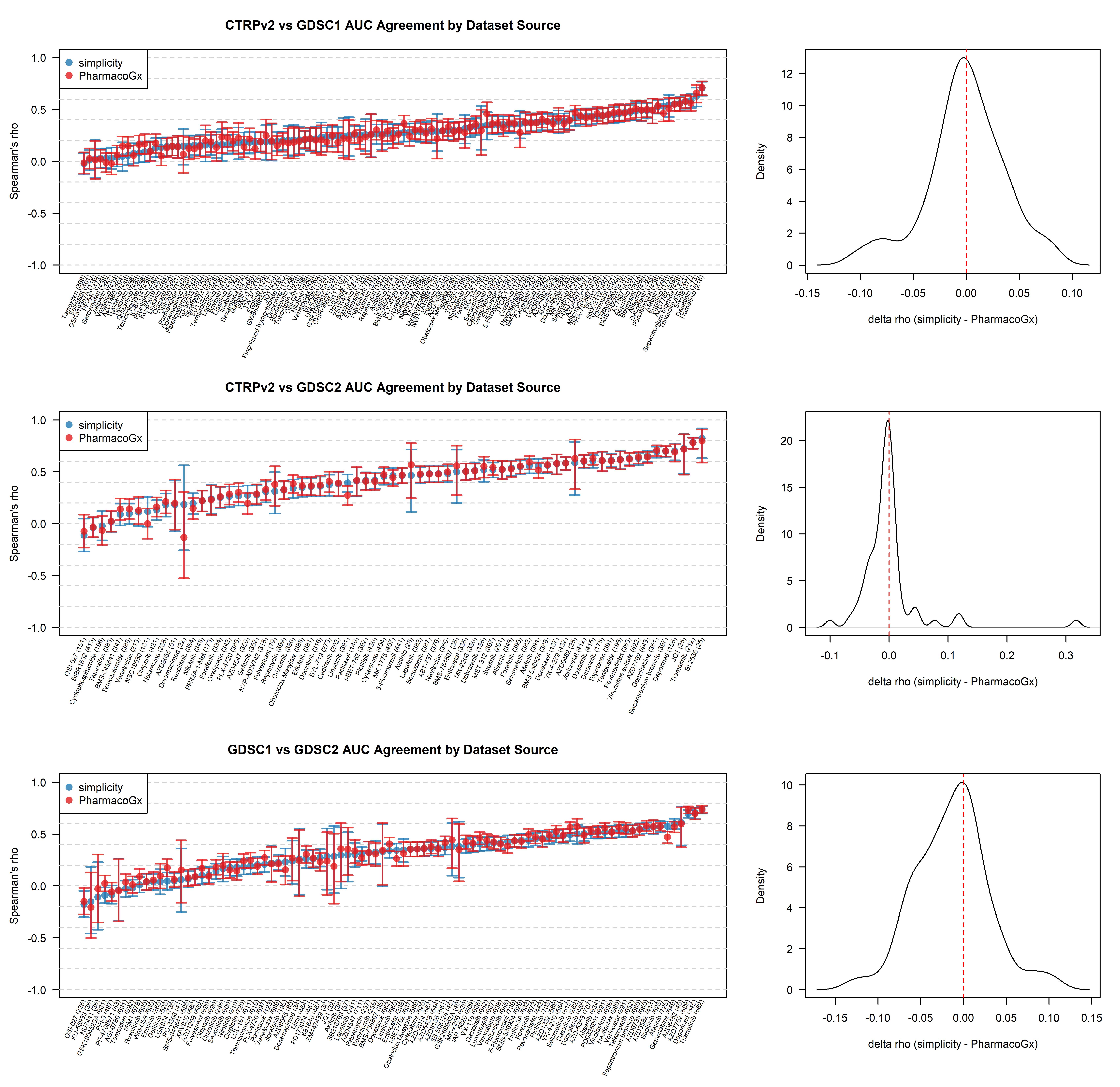
**Figure S2. Agreement between screening datasets using data from Simplicity vs. from PharmacoGx**^9^**.** Agreement between three dataset comparisons is shown: 1. (**A**) CTRPv2 vs. GDSC1; 2. (**B**) CTRPv2 vs. GDSC2; 3. (**C**) GDSC1 vs GDSC2. Agreement between datasets is quantified as Spearman’s rho for AUC values calculated for each compound using overlapping cell lines and drug concentrations. Plots on the **LEFT** show these rho values for each drug with error bars indicating 95% confidence intervals. Density plots on the **RIGHT** show the distribution of differences in rho values between Simplicity and PharmacoGx, with negative values indicating higher rho values using data from PharmacoGx and positive values indicating higher rho values using Simplicity. Parentheses after compound names indicate the number of shared cell lines between the two datasets. Only compounds tested in at least 20 shared cell lines between the datasets were included in the analysis.

**C)**

**B)**

**A)**

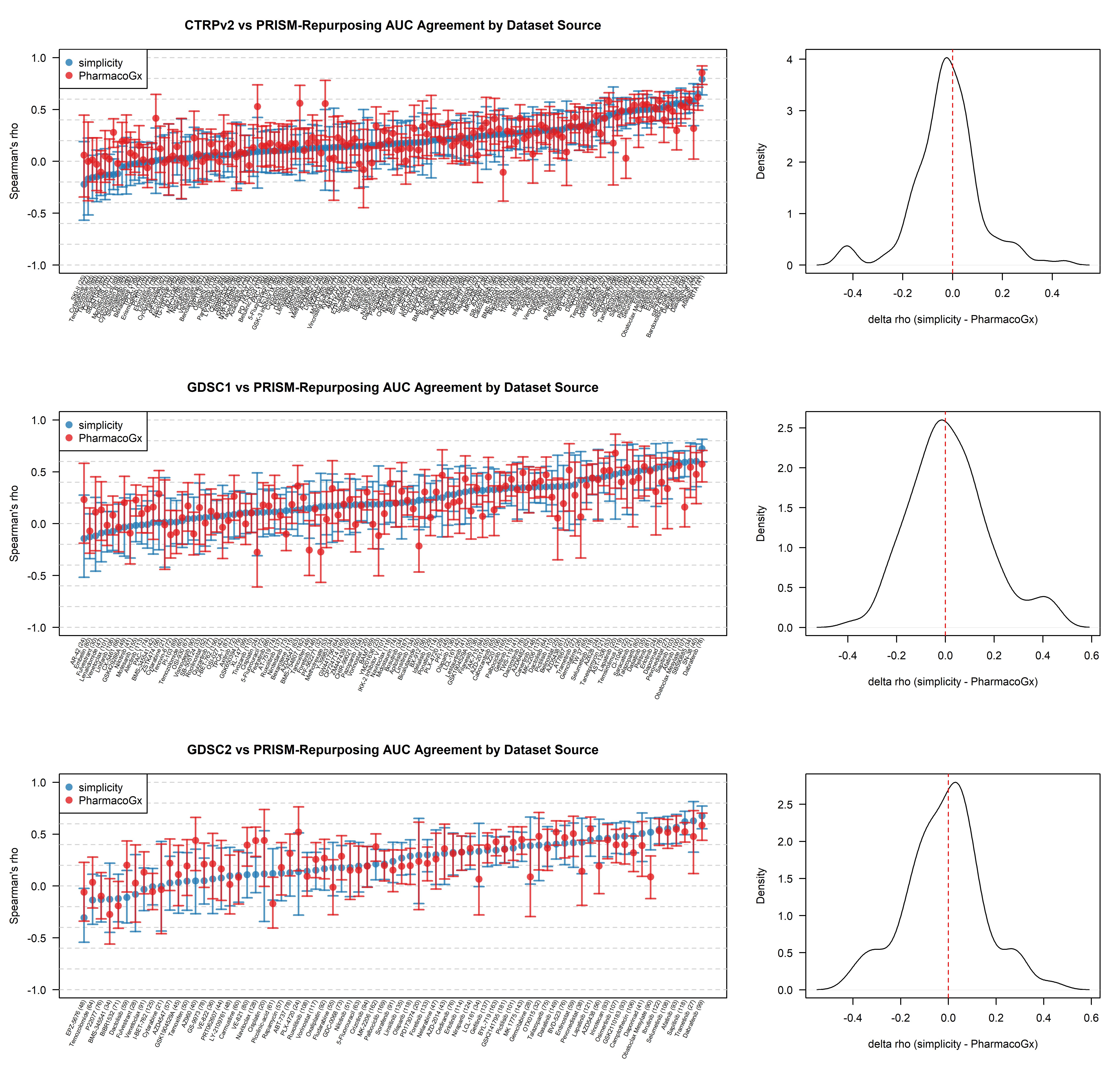
**Figure S3. Agreement between screening datasets using data from Simplicity vs. from PharmacoGx**^9^**.** Agreement between three dataset comparisons is shown: 1. (**A**) CTRPv2 vs. PRISM-Repurposing; 2. (**B**) GDSC1 vs PRISM-Repurposing; 3. (**C**) GDSC2 vs PRISM-Repurposing. Agreement between datasets is quantified as Spearman’s rho for AUC values calculated for each compound using overlapping cell lines and drug concentrations. Plots on the **LEFT** show these rho values for each drug with error bars indicating 95% confidence intervals. Density plots on the **RIGHT** show the distribution of differences in rho values between Simplicity and PharmacoGx, with negative values indicating higher rho values using data from PharmacoGx and positive values indicating higher rho values using Simplicity. Parentheses after compound names indicate the number of shared cell lines between the two datasets. Only compounds tested in at least 20 shared cell lines between the datasets were included in the analysis.

**A)**

**C)**

**B)**

**Figure S4. Top 8 compounds which had higher Spearman’s rho values in PharmacoGx than Simplicity when comparing CTRPv2 vs. PRISM-Repurposing.** Each grouping of 2 plots shows AUC values from PRISM-Repurposing (y-axis) vs. AUC values from CTRPv2 (x-axis). Plots on **LEFT** use data from Simplicity. Plots on **RIGHT** use data from PharmacoGx. AUC values are calculated using only the shared cell lines and concentration range tested between CTRPv2 and PRISM-Repurposing. Plots are shown for the top 8 compounds in which PharmacoGx had higher Spearman’s rho values than Simplicity in Figure S3A.

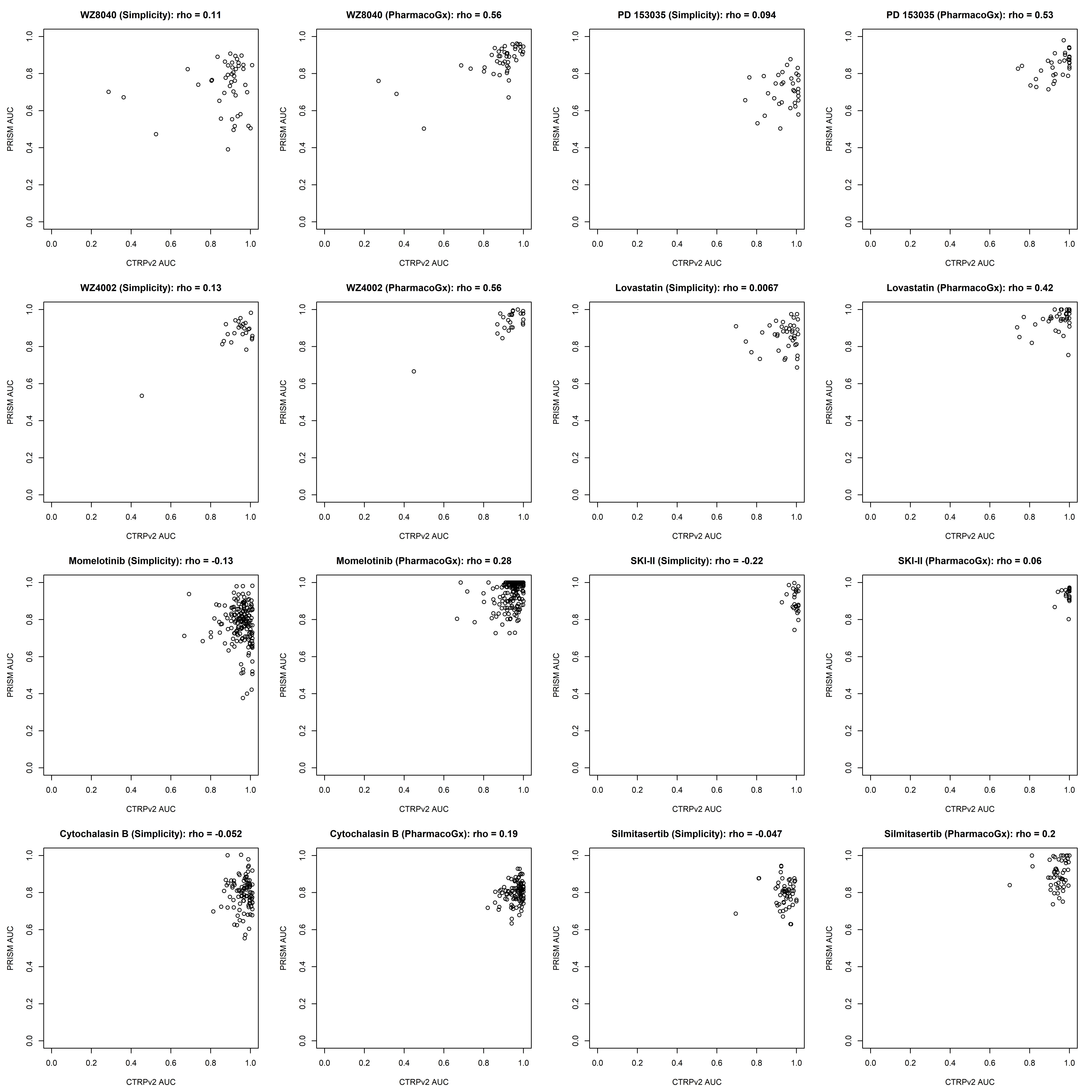

**Figure S5. Top 8 compounds which had higher Spearman’s rho values in Simplicity than PharmacoGx when comparing CTRPv2 vs. PRISM-Repurposing.** Each grouping of 2 plots shows AUC values from PRISM-Repurposing (y-axis) vs. AUC values from CTRPv2 (x-axis). Plots on **LEFT** use data from Simplicity. Plots on **RIGHT** use data from PharmacoGx. AUC values are calculated using only the shared cell lines and concentration range tested between CTRPv2 and PRISM-Repurposing. Plots are shown for the top 8 compounds in which Simplicity had higher Spearman’s rho values than PharmacoGx in Figure S3A.

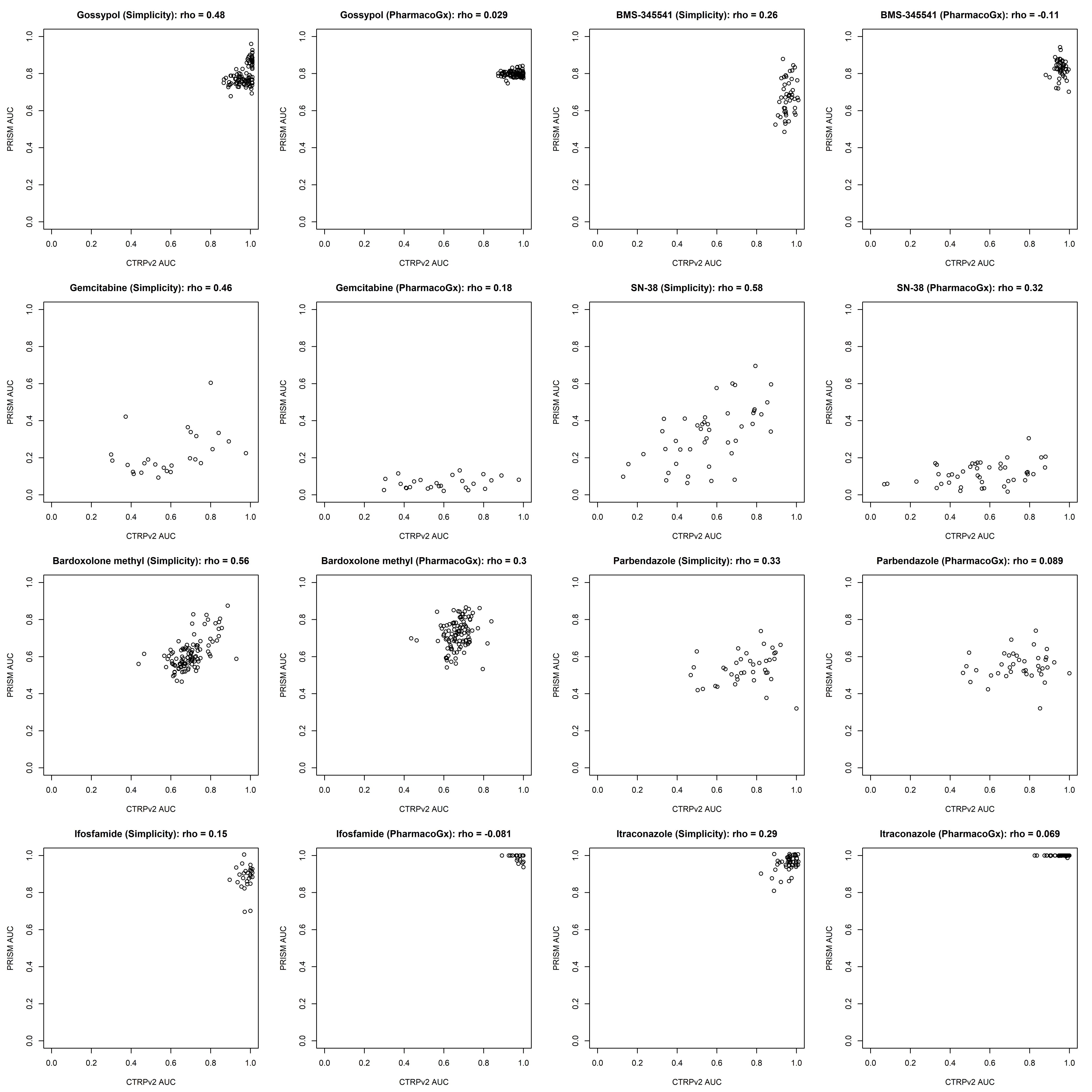
